# SENgEAR: an interactive visualization platform for comparative analysis of senescence signatures and cellular communication

**DOI:** 10.64898/2026.10.31.763339

**Authors:** Sushma Nagaraj, Ricky S. Adkins, Alexandra N. Rindone, Arun Das, Daniel Lesperance, Joshua Orvis, Kavita Krishnan, Benjamin D. Cosgrove, Nathan Basisty, Darren J. Baker, Owen R. White, Anup A. Mahurkar, Jennifer H. Elisseeff, Elana J. Fertig

**Affiliations:** Institute for Genome Sciences, University of Maryland School of Medicine, Baltimore, MD, USA; Translational Therapeutics & Regenerative Engineering Center, Department of Chemical and Biomolecular Engineering and Biomedical Engineering, Johns Hopkins University, Baltimore, MD, USA; Department of Epidemiology, University of Maryland School of Medicine, Baltimore, MD, USA; Department of Medicine, University of Maryland School of Medicine, Baltimore, MD, USA; Greenebaum Comprehensive Cancer Center, University of Maryland School of Medicine, Baltimore, MD, USA; Genetics, Genomics and Development Graduate Program, Cornell University, Ithaca, NY, USA; Meinig School of Biomedical Engineering, Cornell University, Ithaca, NY, USA; Translational Gerontology Branch, National Institute on Aging, NIH, Baltimore, Maryland, USA; Robert and Arlene Kogod Center on Aging, Mayo Clinic, Rochester, MN, USA; Department of Biochemistry and Molecular Biology, Mayo Clinic, Rochester, MN, USA; Department of Pediatrics, Mayo Clinic, Rochester, MN, USA; Paul F. Glenn Center for the Biology of Aging Research, Mayo Clinic, Rochester, MN, USA

## Abstract

Single-cell and spatial transcriptomic datasets provide tremendous potential to identify the cellular states and communication networks associated with senescence. Annotating senescent cells in these datasets is hindered by the lack of universal markers for senescent cells and their context-dependent variability. To overcome this, several groups have developed gene signatures to annotate cells with distinct senotypes. However, comparing these signatures and their biological contexts to determine the specificity of senescence signatures remains complicated by the separate data platforms for signatures and datasets. We present SENgEAR, a no-code web platform for senescent cell annotation, visualization of cell-cell communication networks, and comparison of senescence signatures across studies, tissues, species, and data modalities. SENgEAR provides a unified framework to explore contextual signature specificity, cellular communication pathways, and senotype heterogeneity. We demonstrate the use of SENgEAR for applications spanning aging, fibrosis, and wound healing and its ability to interrelate these states.

## Introduction

Senescent cells are implicated in numerous pathologies related to aging, fibrosis, and cancer, yet it remains challenging to identify them. Cellular senescence has traditionally been recognized as a stress-induced cell state characterized by stable cell cycle arrest involving upregulation of cell cycle inhibitors such as p21 (*Cdkn1a*) and p16 (*Cdkn2a*), and a senescence-associated secretory phenotype (SASP) by which the cells can modulate the tissue microenvironment. Emerging evidence suggests senescence is not a single, uniform program, but a range of states, or senotypes [1–3], that vary across cell types, senescence inducer, time since senescence entry and biological contexts [4–6]. Due to this heterogeneity, no single universal marker can detect senescent cells in all biological conditions.

Single-cell and spatial transcriptomics (ST) datasets provide the potential to characterize these phenotypes across diverse conditions, but the lack of universal marker genes still challenges these analyses. To identify senescent cells in transcriptomics datasets, studies have developed numerous senescence gene signatures via literature curation [7], *in vitro* perturbations [5, 8], or *in vivo* models [9], ranging from small, unweighted gene lists to weighted lists with hundreds of genes. While these signatures have been validated to identify senescent cells in specific contexts, it is unknown how these signatures relate to senescence in other cell types, tissue types, and disease states. Once identified, the context of these cellular phenotypes and signaling with other cells in the environment are crucial to their modulation. Therefore, there remains a need to readily perform cross-signature and cross-context analyses to better understand which senotypes these signatures represent, their conservation across species and biological contexts and their influence on tissue state via their SASP.

## Results

### SENgEAR enables senescence analysis with automated transfer learning and visualization of cell-cell communication networks

To enable exploration of senescent cells and their cellular communication networks across biological contexts, we present SENgEAR, a web platform designed to enable systematic annotation of senescent cells across heterogeneous datasets and exploration of cell-cell communication (https://senescence.umgear.org). Built on the gEAR [10] ecosystem, it allows for point-and-click multiomics data visualization, with results shared through permalinks to facilitate collaboration and reproducibility. The platform supports analysis of a wide variety of datatypes (**Figure 1A**) and its extension to senescence in SENgEAR hosts curated collections of single-cell and ST datasets spanning aging, fibrosis, and cancer (**Figure 1B**, **Supplementary Table 1**). Users can upload their own datasets and interactively visualize gene expression across uploaded and curated data collections (**Figure 1C**).

**Figure 1.**
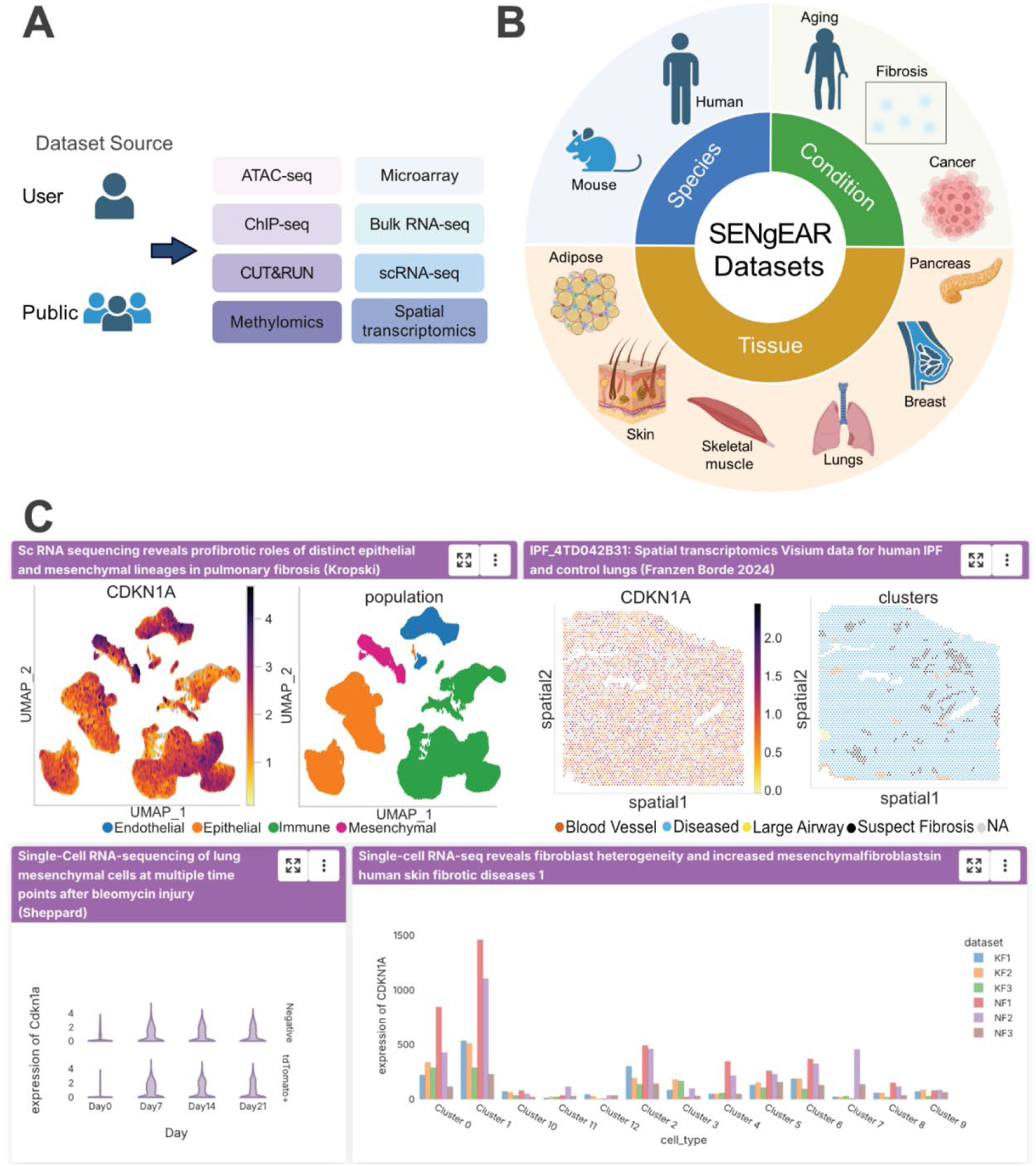
Overview of SENgEAR platform on https://senescence.umgear.org. **A)** Omics datatypes compatible with SENgEAR. Platform enables users to upload their own datasets or datasets from publicly available sources. **B)** Species, tissues, and biological conditions represented by datasets in SENgEAR. **C)** Representative data visualizations available in SENgEAR. Platform enables users to visualize different datasets and plot types simultaneously.

A unique feature of the SENgEAR platform is the automated annotation of transcriptomics datasets from gene signatures using transfer learning [11] (**Supplementary Figure 1**), allowing users to compare senescence signatures across studies. To facilitate this analysis, SENgEAR incorporates a diverse collection of published senescence signatures capturing several biological contexts and derivation strategies (**Figure 2A-B**, **Supplementary Table 2**), including signatures associated with aging, cancer, fibrosis, and wound healing. Automated ortholog mapping enables direct comparison of human and mouse datasets, allowing users to evaluate the conservation and context dependence of senescence signatures across species.

**Figure 2.**
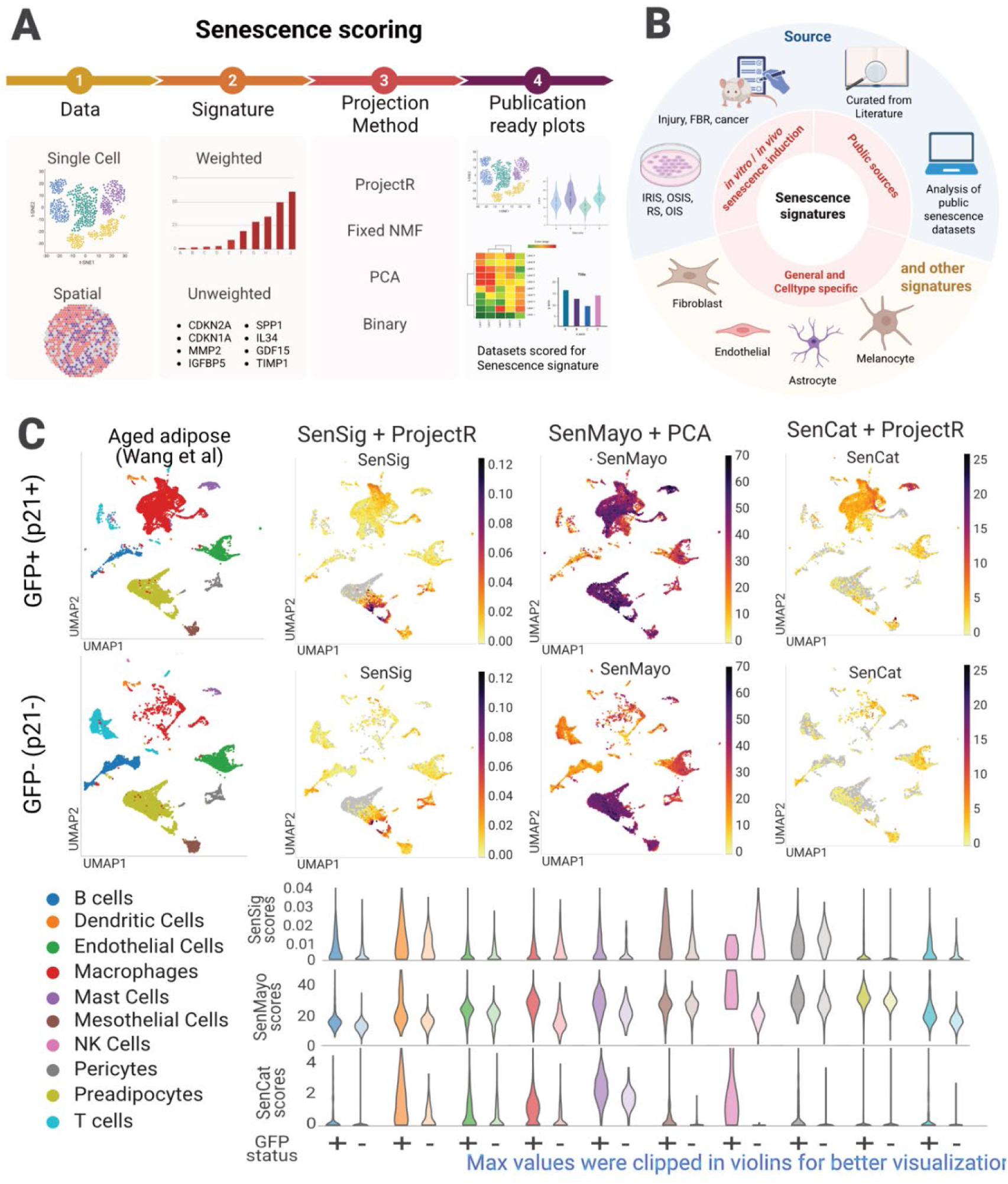
Cross-comparison of senescence signatures using SENgEAR. Overview of **A)** projection analysis workflow and **B)** senescence signatures integrated into SENgEAR. **C)** Comparison of signature projections in experimentally defined adipose cell populations from p21-Cre-GFP mice [12], demonstrating differences in the specificity of senescence signatures.

### Comparison of senescence signatures with SENgEAR

To demonstrate cross-signature comparisons in SENgEAR, we first analyzed a single-cell RNA-seq (scRNAseq) dataset of sorted p21-Cre-GFP-labeled adipose cells from aged mice [12] (**Figure 2C**). This dataset provides an experimentally defined cell population enriched for senescence-associated features [12, 13], enabling evaluation of how different senescence signatures classify cells in a well-characterized biological context. For the cross-comparison, we selected three distinct signatures derived from different biological conditions to provide a representative sampling of the diverse senescence signatures in SENgEAR. We performed transfer learning on the dataset with SenSig [9, 14], an *in vivo*-derived signature of p16+ senescent stromal cells; SenCat [8], a signature of core senescence markers based on several *in vitro* senescence induction experiments; and SenMayo [7] a widely used literature curated gene set that has been validated in aging contexts. This analysis revealed differences in the specificity of these signatures to p21-high cells in aged adipose tissue. SenCat displayed selective enrichment and higher scores in p21-high immune cell populations, whereas SenSig and SenMayo appeared in both p21-high and p21-low cell populations (**Figure 2C**). Our observations are consistent with the concept that different senescence signatures capture distinct senotypes, and that their specificity to senescent cells is dependent on biological context.

### Cell to cell communication analysis

Senescent cells influence their microenvironment by secreting SASP factors and expressing cell surface molecules that bind to cognate interacting partners in neighboring cells. These binding interactions form important cell to cell communication networks that impact the tissue microenvironment. To explore this broad impact of senescent cells, we extended the SENgEAR platform to enable users to visualize and explore cell-cell communication networks inferred from ligand-receptor-transcription factor cellular network inference tool dominoSignal [15, 16] (**Supplementary Figure 2**). Briefly, dominoSignal infers cellular signaling pathways by linking ligand-receptor interactions to transcription factor activity in receiver cell populations (Methods). This approach enables the prediction of biologically meaningful cell-cell signaling interactions that activate downstream pathways in receiver cells. However, visualization of the numerous molecular and cellular components underlying the inferred signaling networks has remained a challenge. Therefore, we developed a new cell to cell communication module within the gEAR [10] software to facilaite visualization of these interactions.

To demonstrate the application of SENgEAR to study these signaling networks and interactions, we first performed dominoSignal cell-cell communication inference on the p21-Cre-GFP-labeled adipose dataset [12]. To interpret the dominoSignal results, we utilized the cell-cell communication interface implemented in SENgEAR (**Figure 3A**). Through this interface, we found that p21-high macrophages, the most prevalent p21-high cell population in this dataset, engaged in more inflammatory signaling with immune cell populations compared to p21-low macrophages (**Figure 3B**). In particular, these p21-high macrophages expressed pro-inflammatory cytokines, such as *Ccl2*, *Ccl3*, and *Ccl7*, that bind to CC chemokine receptors in several of the innate and adaptive immune cell populations. To predict how these signaling interactions affect transcriptional changes in receiver immune cells, we identified transcription factors that correlated with expression of *Ccr1*, a receptor that binds multiple CC chemokines (**Figure 3C**). These include transcription factors involved in inflammatory pathways and immune cell activation, such as Nf-κB activation (*Nfkb1*, *Nfkb2*) and interferon response (*Stat1*, *Irf2*, *Irf8*). These data corroborate previous findings that p21+ cell elimination reduces expression of such inflammatory pathways in murine visceral adipose tissue [12]. Taken together, these analyses illustrate how SENgEAR can be applied to visualize senescence-specific cell-cell signaling interactions that inform mechanistic studies.

**Figure 3.**
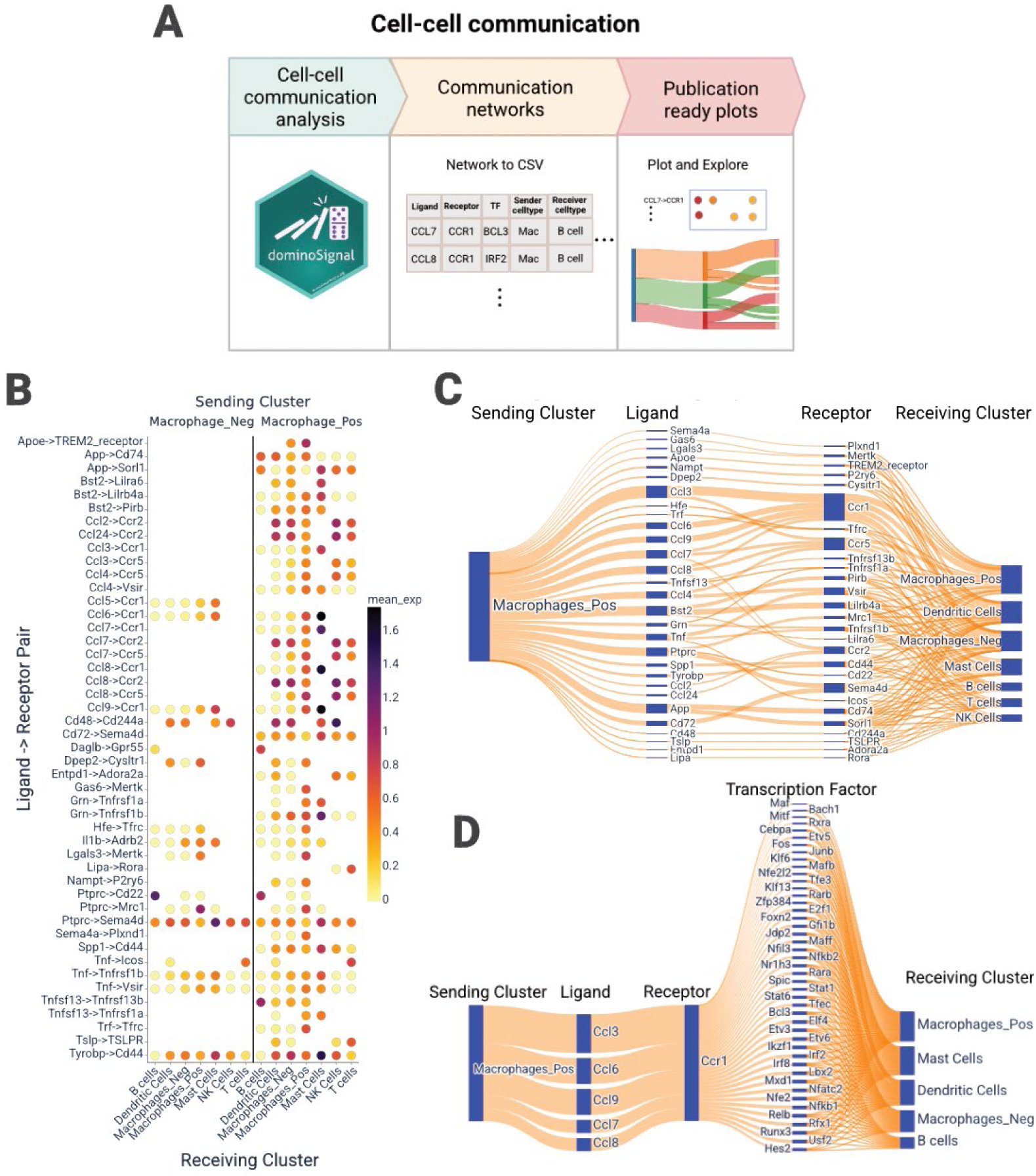
Cell-cell communication in SENgEAR. **A)** Workflow to assess cell-cell communication of scRNAseq data in SENgEAR. Steps include performing cell-cell communication analysis using packages such as dominoSignal [15, 16], converting the signaling network to a CSV file, and plotting and exploring data in the SENgEAR portal. **B-D)** Cell-cell communication between p21-high or p21-low macrophages (sender) and immune cells (receiver) in aged murine adipose tissue [12]. Dot plot in **B** shows the geometric mean of average z-scored ligand receptor expression by sending clusters and average z-scored receptor expression by receiving clusters. River (sankey) plots in **C** and **D** show the signaling network between ligands expressed by p21-high macrophages and receptors expressed by receiver immune cells. Plot in **D** shows transcription factors that are correlated with *Ccr1* receptor expression in receiver immune cells.

### Cross-species and cross-modality analysis

To assess the utility of SENgEAR to perform cross-context comparisons across tissues and species, we analyzed scRNA-seq datasets from a mouse model of foreign body response [17] (FBR) and from human idiopathic pulmonary fibrosis [18, 19] (IPF). These datasets represent two different fibrotic conditions that have been previously linked to senescence [20, 21]. By projecting these datasets into SenSig, SenMayo, and SenCat, we found that each signature exhibited similar expression patterns across the two datasets (**Figure 4A-B**). SenMayo scores were high in both myeloid cells and fibroblasts, whereas SenCat scores were high in myeloid cells and SenSig scores were high in fibroblasts. In addition, while elevated SenMayo and SenCat scores were observed more broadly across different cell populations in each dataset, SenSig was primarily restricted to fibroblast cell populations that expressed higher levels of *Cdkn2a* (p16; **Supplementary Figure 3**). This finding is consistent with other studies showing that p16+ senescent fibroblasts are present in IPF and the FBR [9, 21], suggesting that SenSig may capture a conserved p16-associated senescence phenotype across different fibrotic pathologies.

**Figure 4.**
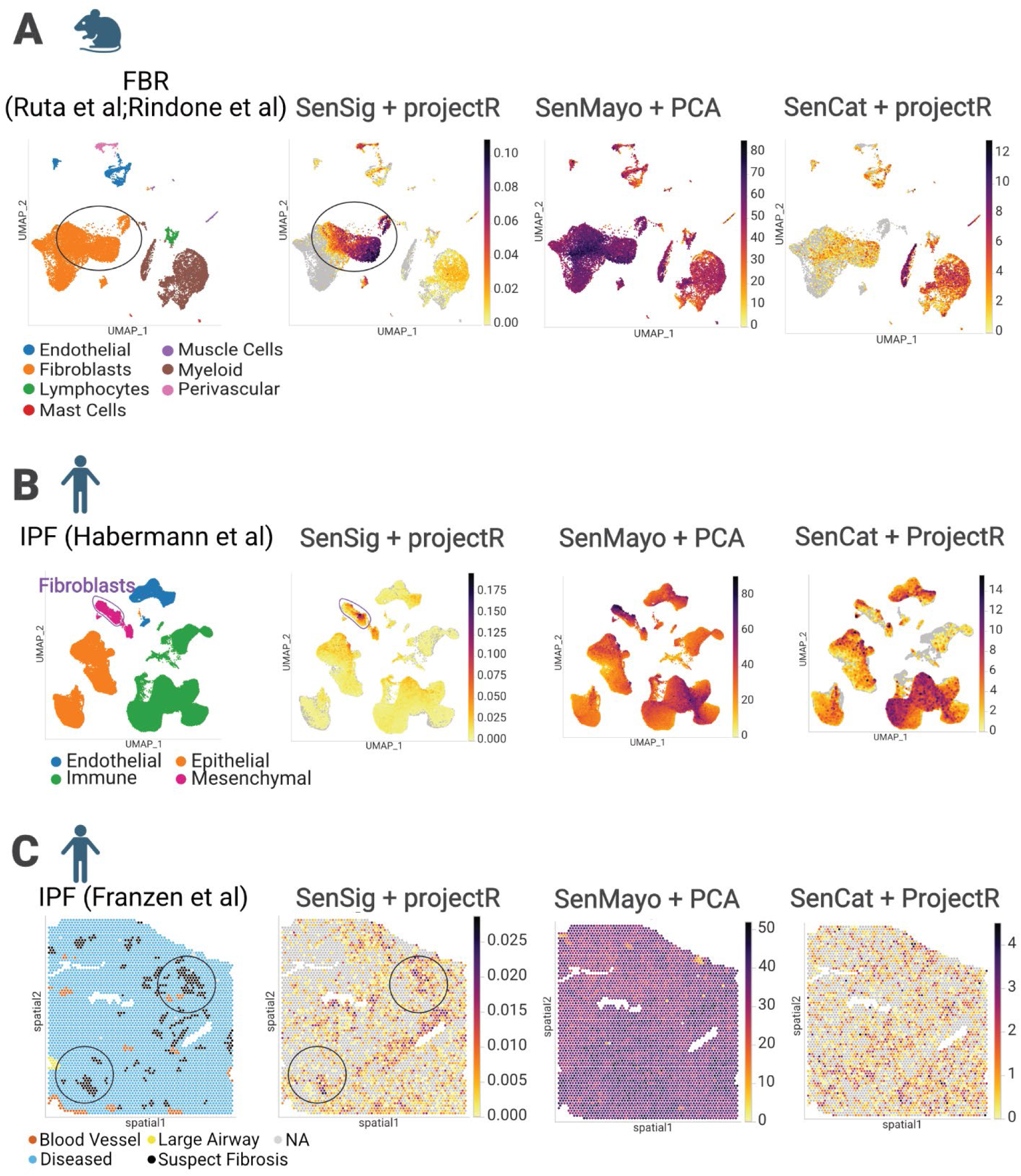
Comparison of senescence signatures in fibrotic conditions across species and datatypes. Datasets shown represent **A)** scRNAseq of murine fibrosis resulting from foreign body response (FBR) to a biomaterial implant [14, 17], **B)** scRNAseq of human idiopathic pulmonary fibrosis (IPF) [18], and **C)** Visium of human IPF [19]. Black circles highlight cell fibroblast/mesenchymal cell populations with a high SenSig projection weight.

To further demonstrate SENgEAR’s utility for cross-platform analysis, we also analyzed these gene signatures in human IPF ST data [19]. Spots with elevated SenMayo and SenCat scores were present throughout the diseased tissue and lacked regional specificity. By contrast, SenSig enrichment was localized to regions associated with fibrosis, suggesting that the senescent fibroblasts we identified using scRNA-seq reside in the fibrotic regions of the IPF tissue (**Figure 4C**). To further probe the phenotype of these senescent fibroblasts, we projected the ST data into separate NMF-derived fibroblast senotype patterns previously learned from the mouse FBR dataset [14, 17] (**Figure 5**). These include patterns related to cell cycle arrest (pan-senescent fibroblast pattern), immune signaling, cartilage development, cell adhesion and signaling, and fibrosis. We previously showed that these patterns are expressed by different fibroblast senotypes that localize to distinct spatial niches in the FBR [14]. By applying these patterns to the ST IPF dataset, we found that SenSig-enriched spots had higher scores for the pan-senescence cell cycle arrest pattern and the cartilage development pattern, a pattern expressed by a fibroblast senotype associated with mature fibrotic ECM [14] (**Figure 5A**). By contrast, we did not observe enrichment of any of the senotype patterns in healthy control tissue (**Figure 5B**). Together, these results show that SENgEAR can be used to resolve fine-grained senotypes across tissues and species.

**Figure 5.**
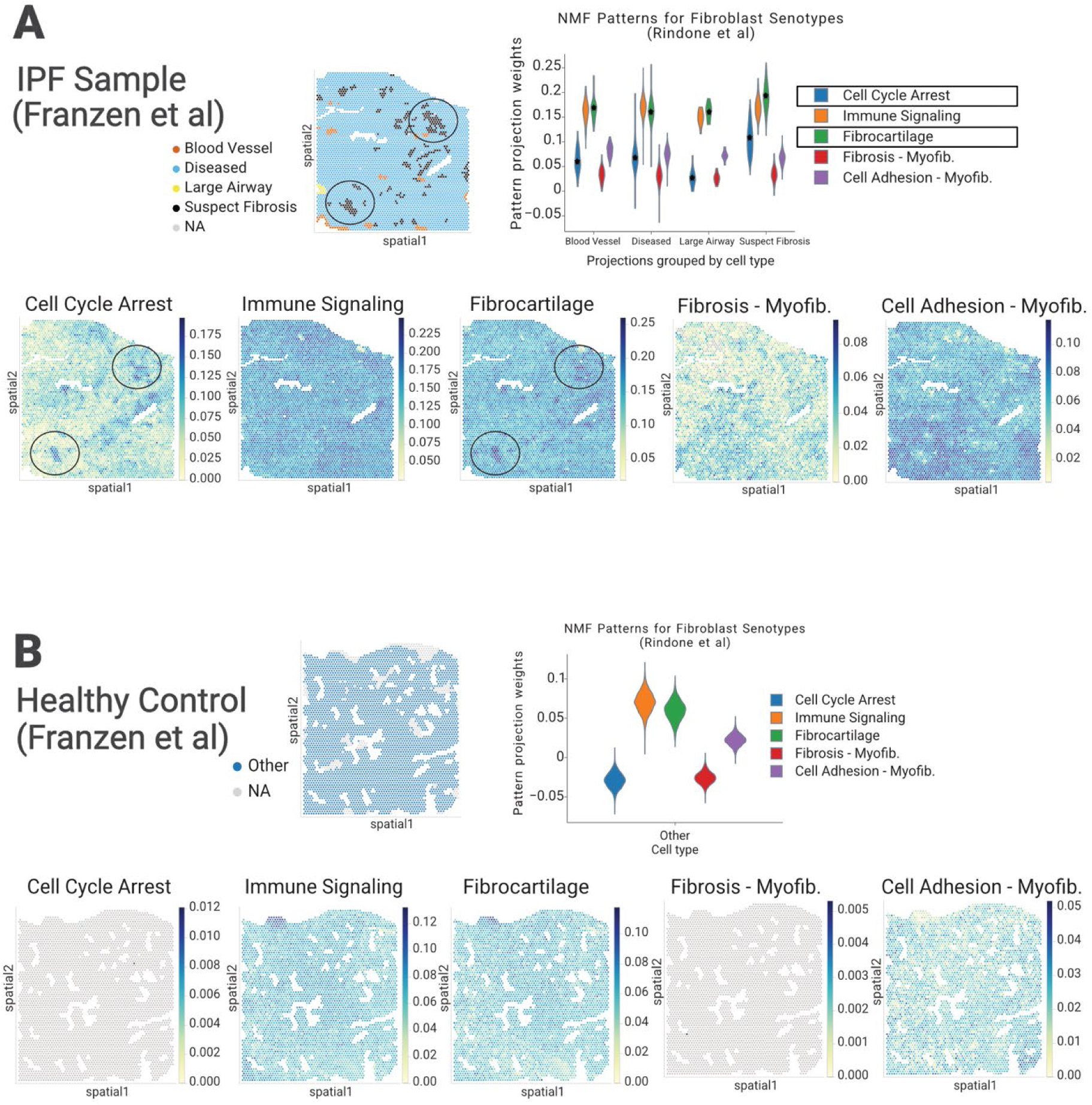
Fibroblast senotypes are localized in fibrotic regions in IPF. Projection of IPF ST data [19] into fibroblast senotype patterns learned on the Mouse FBR single-cell data [14, 17]. **A)** Black circles show SenSig enriched spots in IPF sample that have higher weights for the pan-senescence cell cycle arrest pattern and the fibrocartilage (i.e., cartilage development) senotype pattern [14]. **B)** Healthy sample does not show enrichment for any of the senotype patterns.

### Senescence dynamics during mouse skeletal muscle regeneration with aging

Because the composition and transcriptional features of senescence-associated cell populations can shift over time based on factors such as wound healing and aging, it is important to evaluate the temporal behavior of senescent cells. To demonstrate the utility of SENgEAR to examine senescence dynamics during wound healing across different ages, we applied it to a single-cell dataset of skeletal muscle regeneration [22], a process known to be influenced by age-associated changes in cellular state. We projected the longitudinal single-cell dataset of murine muscle injury across age groups into SenSig [9, 14], which was previously shown to capture senescent-like cell populations in this context [22]. This analysis revealed differences in senescence temporal dynamics between different cell types and ages (**Figure 6**). Senescence scores increased over time in fibro-adipogenic progenitors (FAP) and muscle stem cells (MuSCs) and progenitors, with a peak 3.5 days after injury, consistent with transient senescence induction during muscle healing [23, 24]. By contrast, senescence scores in pericytes and vascular smooth muscle cells peaked at 1 and 2 days following injury in the young and old age groups. In young mice, senescence scores in all cell populations generally declined as regeneration progressed, whereas in geriatric animals, they remained elevated, consistent with impaired resolution and reduced regenerative capacity [22]. These results show that SENgEAR can be used to capture dynamic, age, and cell type-dependent changes in senescence that align with known differences in regenerative outcomes.

**Figure 6.**
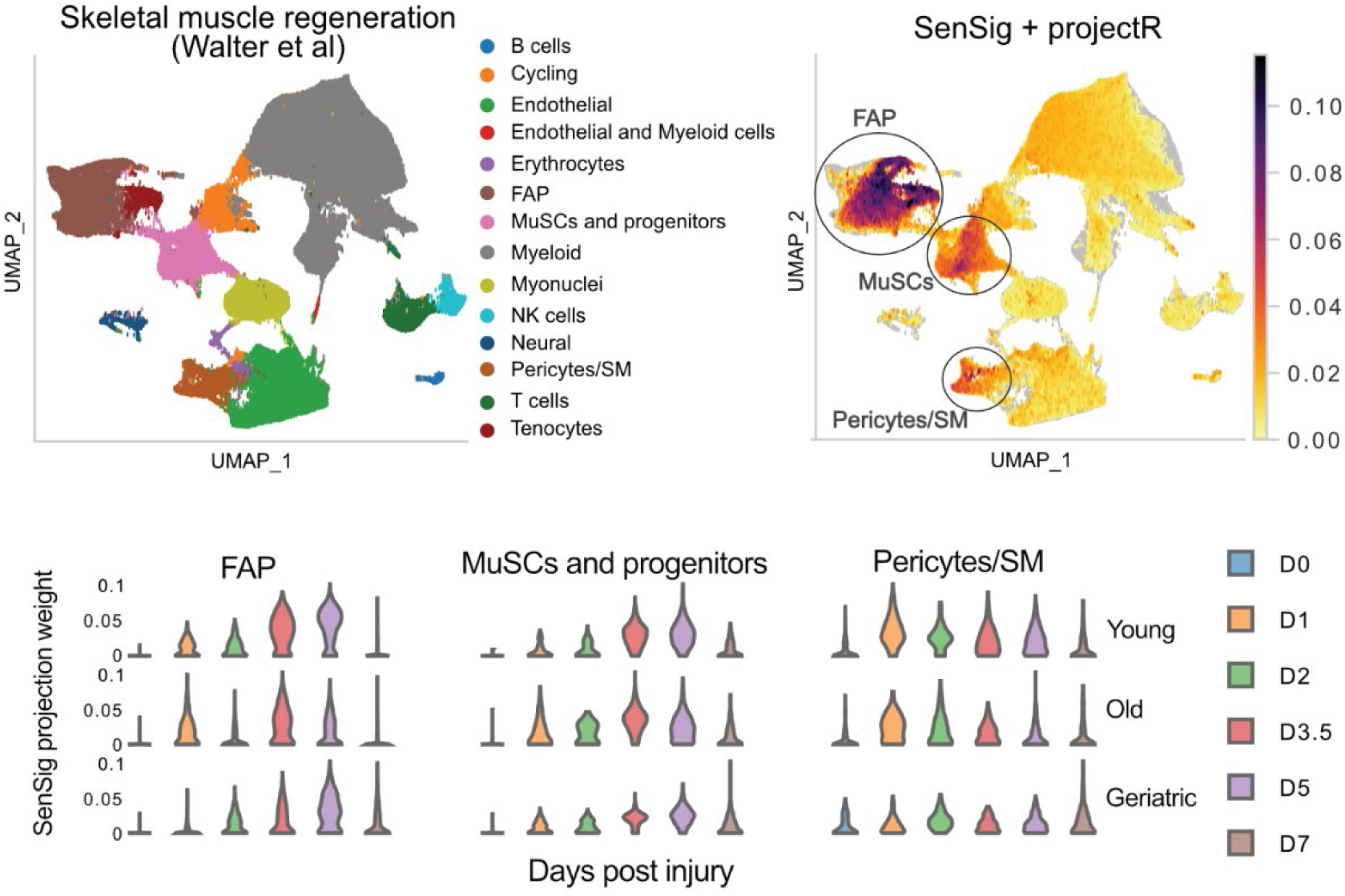
Senescence dynamics during skeletal muscle regeneration in young, aged, and geriatric mice from 0 to 7 days following notexin injury. (Top) Projection of a scRNA-seq atlas of muscle regeneration (Walter et al; left) into SenSig (right) identifies senescence as enriched within the fibro-adipogenic progenitors (FAP), muscle stem and progenitor cells (MuSCs), and pericytes/smooth muscle cells (Pericytes/SM). (Bottom) Violin plots display the dynamics in SenSig projection weights by cell types, ages, and timepoints post-injury.

## Discussion

SENgEAR provides a unified framework for applying, comparing, and interpreting senescence signatures across diverse datasets, experimental systems, and species. Combined with the embedded cell-cell communication visualization interface, it builds a foundation for predicting context-specific regulatory mechanisms by which senescent cells influence their surrounding microenvironment. By integrating multiple analytical approaches, curated datasets, senescence signatures, and automated cross-species mapping within an accessible interface, SENgEAR lowers the technical barrier to senescence analysis providing a resource to the senescence community. While our study focuses on datasets of biological conditions previously linked to senescence, including fibrosis and adipose aging, SENgEAR provides users with the ability to upload gene signatures and datasets, curate dataset collections, and create custom visualizations. This framework will facilitate reproducible cross-comparison studies of senescence across species and biological contexts.

SENgEAR builds upon the gEAR [10] platform by extending functionality to visualization and exploration of cell-cell communication networks in addition to the integrated transfer learning methods. While tailored to study senescence in this instance, the platform and its integrated tools have broad utility in other areas including hearing research [10], neuroscience [25] and cancer research [26]. In addition, the curated format of SENgEAR can be readily adapted to enable rigorous, biologically driven AI queries that accelerate data analysis and discovery. Future work will prioritize integrating AI models and interactive Jupyter notebooks within SENgEAR to support custom downstream analysis.

Cell-cell signaling networks in tissues are critical mediators of senescent cell phenotype and function [3]. The extension of the SENgEAR platform to explore cell-cell communication is vital to characterizing these regulatory networks. While this study used dominoSignal [15, 16] to perform the cell-cell communication analysis, SENgEAR uses a tabular format for data ingest and can be extended to visualize results from other cell-cell communication analysis methods that incorporate the communication between ligands, receptors, and transcription factors [27, 28]. Our format provides a generalized interface and can be readily adapted to support methods to study cell-cell signaling in ST datasets as future areas of extension. Future work must also go beyond pre-computed cellular communication to interactive analyses of the datasets, allowing for refined annotation of senotypes in the datasets from the in-built gene signature analysis into these communication visualizations.

## Methods

### Datasets in SENgEAR

A comprehensive list of datasets in SENgEAR are listed in **Supplementary Table 1**. These datasets represent conditions such as aging, wound healing, fibrosis, and cancer in both mouse and human tissues [9, 12, 17–19, 22, 29–35]. Datasets were uploaded to SENgEAR in a variety of formats including H5AD, MEX format (a three-tab format of gene-level expression, and observation-level tabular files), and Excel spreadsheets. To upload Seurat objects, the metadata, genes, and expression data were extracted into tabular files and uploaded to the portal. Most datasets were converted into H5AD format during the upload process. Datasets generated from spatial assays were transformed into a Zarr store. Aside from the data conversion to H5AD/Zarr store, no further pre-processing or normalization was performed during the upload process.

### Senescence signatures in SENgEAR

A comprehensive list of signatures in SENgEAR is provided in **Supplementary Table 2** [5, 7–9, 14, 23, 36–50]. All the signatures were obtained from previously published articles as provided in the supplemental texts by the original authors or from MSigDB [51, 52]. The gene signatures were formatted for SENgEAR and uploaded to the portal.

### Projection analysis

All the projection analyses were performed using the projection methods and analysis tools integrated into SENgEAR. For projection into weighted signatures, we used the embedded transfer learning method projectR [11] which implements a least-squares fit of the signature to the target dataset. For projection with unweighted signatures we used the SENgEAR implementation of PCA projection with a constant weight of 1 assigned to all the genes in the signature. This method computes a dot product of dataset gene expression and of the gene signature weights and can be used as a general-purpose method with both weighted and unweighted signatures. Additionally, an NMF algorithm as implemented in SJD [53] is available where the gene weights matrix (signatures) from a prior NMF run can be provided to estimate new sample weights. SENgEAR also includes a simple binary count for unweighted gene lists which counts the number of genes from the gene list with non-zero expression levels in the target dataset.

Different projection methods were applied based on the format of the senescence signature (**Supplementary Table 2, 3**). Projection into SenSig [9, 14] was performed using projectR (selected algorithm Least-squares optimization for NMF) with z-score gene expression normalization, as described previously [14]. Projection into SenMayo [7] was performed using PCA projection (selected algorithm Principal component analysis). Projection into SenCat [8] and NMF signatures from Rindone et al. [14] was performed using projectR (selected algorithm Least-squares optimization for NMF). For all projections, negative coefficient weights were set to zero to facilitate better visualization.

### Cell-cell communication analysis

The R package dominoSignal [15, 16] (version 1.0.5) was used to perform cell-cell signaling analysis. The CellPhoneDB [54] database (version 4.0.0) was used as the interaction database for the dominoSignal analysis. The cells were first scored for transcription factor activity using the Python package SCENIC [55], [version 0.12.1] and dominoSignal was used to infer signaling networks in the data, prioritizing interactions where the receptor expression was correlated with transcription factor activity. dominoSignal was run using the parameters use_clusters = T, use_complexes = T, remove_rec_dropout = F, min_tf_pval = .001, max_tf_per_clust = Inf, max_rec_per_tf = Inf, rec_tf_cor_threshold = .25. The cell-cell signaling networks in the resulting dominoSignal object were converted to a table and saved as a csv file for upload into SENgEAR.

For a given DominoSignal output, the cell-cell communication visualization tool in SENgEAR first converted the tabular csv file into a .parquet file to lower computational overhead. The .parquet file was then processed to generate the following plots: 1) dot plots showing the geometric mean of the average z-scored ligand and receptor expression across specified ligand-receptor and sending-receiving cluster pairs and 2) river plots (also referred to as Sankey diagrams) showing the global network between ligands of sending clusters, receptors of receiving clusters, and transcription factors correlated with receptors of receiving clusters. Both plots were generated using the Python Plotly package (version 6.6.0, https://github.com/plotly/plotly.py/blob/main/CITATION.cff). For both types of visualizations, select plots were filtered for specific cell populations, ligands, and/or receptors using functions built into SENgEAR.

### Data visualization

The data visualizations used for Figures in this study were created using the Single-gene (https://senescence.umgear.org/dataset_curator.html) and Multi-gene (https://senescence.umgear.org/multigene_curator.html) displays on the SENgEAR portal. The new cell-cell communiation visualizations are created using the Cell-cell display on the SENgEAR portal (https://senescence.umgear.org/cell_cell.html). Links to the visualizations for each analysis are provided in **Supplementary File 1**.

## Data availability

The datasets and signatures can be accessed on the SENgEAR portal at https://senescence.umgear.org/dataset_explorer.html and https://senescence.umgear.org/gene_list_manager.html respectively.

## Code availability

R code used to reformat the signatures for upload to SENgEAR is available at https://github.com/FertigLab/SENgEAR-paper. Code for the gEAR platform is available at https://github.com/IGS/gEAR.

## AI usage

Generative AI with Microsoft Copilot was used to assist in writing, supporting initial outlining of the manuscript and improving phrasing with Copilot and Grammarly. Portions of the SENgEAR code base and cell to cell communication extensions were developed with Claude.

## Acknowledgements

We would like to thank Chris Cherry for feedback and support in the development of the original analysis pipelines that led to implementation in the SENgEAR platform. This study was supported by NIH grants U54AG079779 (E.J.F., J.H.E., B.D.C., D.J.B.) and U54CA274371 (E.J.F.). This research was supported in part by the Intramural Research Program of the National Institutes of Health (NIH). The contributions of the NIH author(s) were made as part of their official duties as NIH federal employees, are in compliance with agency policy requirements and are considered works of the United States Government. However, the findings and conclusions presented in this paper are those of the author(s) and do not necessarily reflect the views of the NIH or the U.S. Department of Health and Human Services. This research was also supported in part by University of Maryland, Baltimore through a grant from the Pivot Strategic Investment Initiative. The content is solely the responsibility of the authors and does not necessarily represent the official views of the University of Maryland, Baltimore. Select schematics were created using Biorender: Nagaraj, S. (2026) https://BioRender.com/sqk458t.

---

**Supplementary Figure 1.**
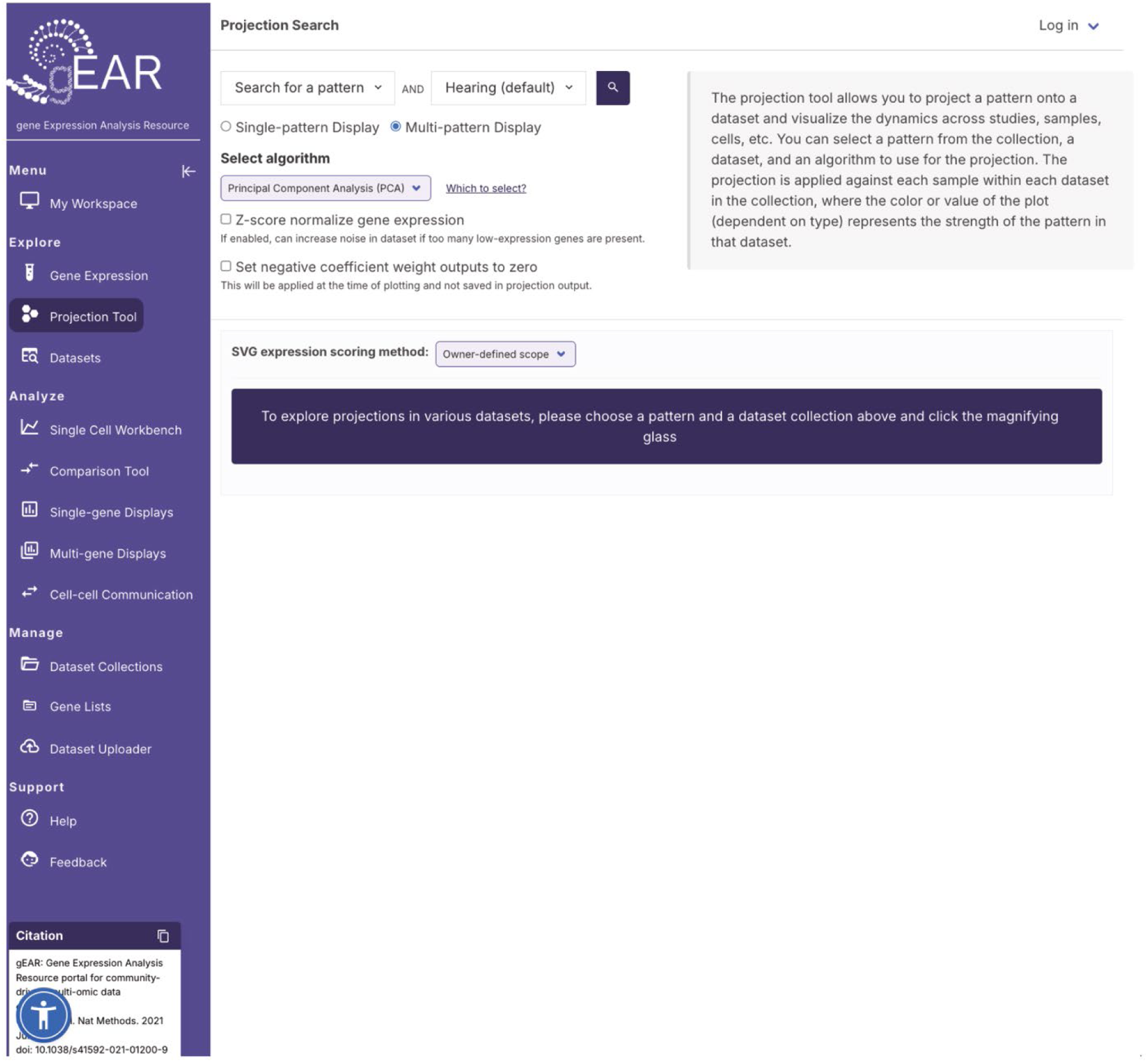
Projection tool on the gEAR platform.

**Supplementary Figure 2.**
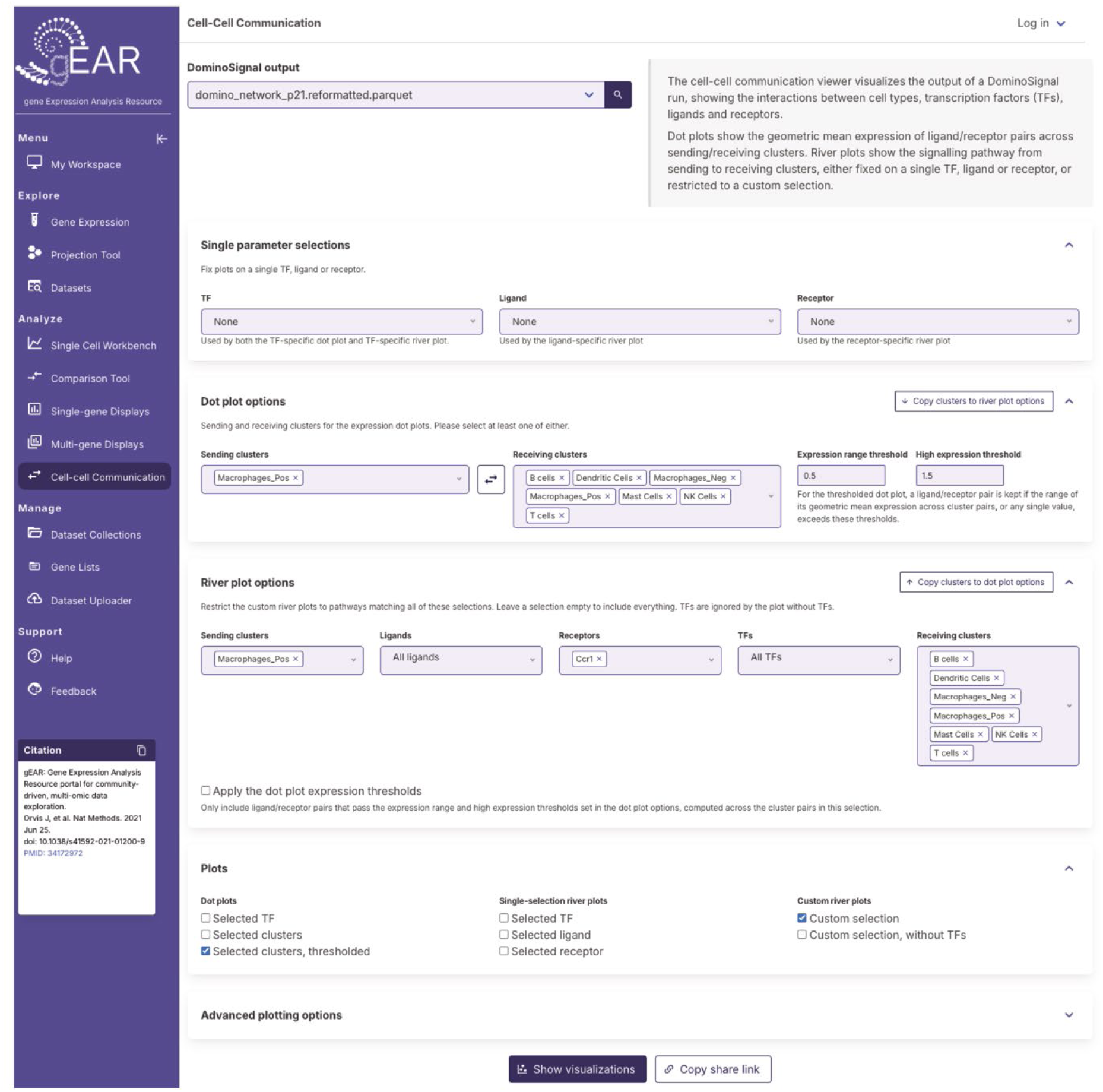
Cell-cell communication visualization and exploration on the gEAR platform.

**Supplementary Figure 3.**
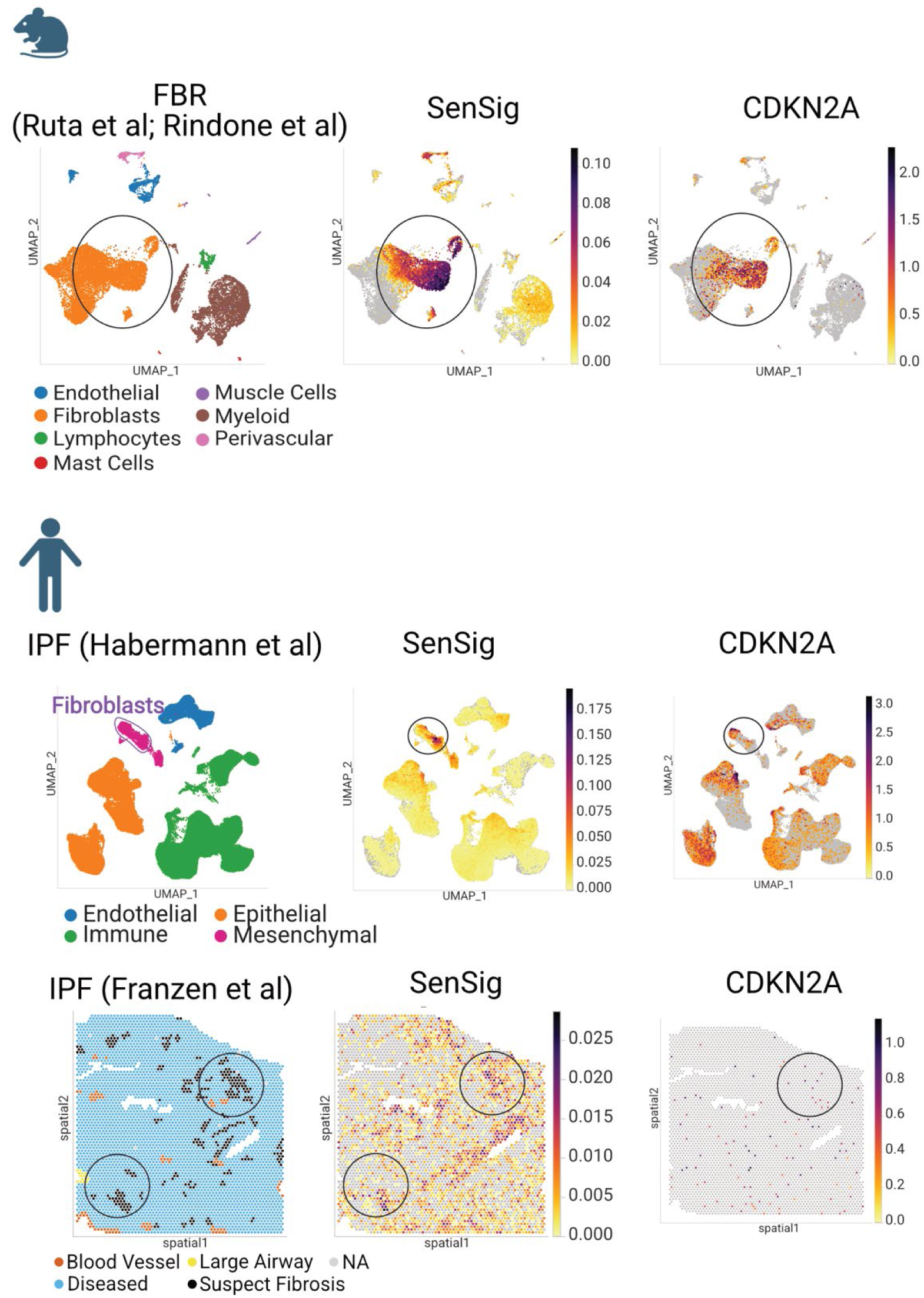
SenSig projection weights are highest in fibroblast populations that express *Cdkn2a* (p16) in Mouse FBR datasets [14, 17] (top) and Human IPF [18, 19] (middle, bottom). Black circles highlight SenSig-high cell populations that express *CDKN2A/Cdkn2a* (p16). Although the expression of CDKN2A is sparse, select SenSig-high spots and *CDKN2A*+ spots overlap with the Suspect Fibrosis regions in the IPF ST dataset [19] (bottom).

**Supplementary Table 1.** List of datasets in SENgEAR.

| Signature Name | Species | Weighted | # Signatures | Method | Cell/Tissue Type | Reference |
|---|---|---|---|---|---|---|
| SenCID | Human | Yes | 6 | Machine learning from published datasets | Multiple | Tao et al., Cell Metab, 2024 |
| p21 PASP | Human | No | 1 | in vitro | Fibroblast | Sturmlechner et al., Science, 2021 |
| SenMayo | Human | No | 1 | Curated from literature | N/A | Saul et al., Nature Commun, 2022 |
| FRIDMAN_SENESCENCE_ (UP, DOWN) | Human | No | 2 | Curated from literature | N/A | Fridman and Tainsky, Oncogene, 2008 |
| GOBP_CELLULAR_SENESCENCE | Human | No | 1 | Curated from literature | N/A | The Gene Ontology Consortium |
| REACTOME_CELLULAR_SENESCENCE | Human | No | 1 | Curated from literature | N/A | Orlic-Milacic, M, Reactome<br>Jupe et al., J Thromb Haemost. 2012 |
| Hernandez-Segura | Human | Yes and No | 7 and 4 | in vitro | Fibroblast, Keratinocytes, Melanocytes, Astrocytes | Hernandez-Segura et al., Curr Biol, 2017 |
| Moiseeva | Mouse | Yes | 12 | in vivo | Myeloid cells, muscle satellite cells (SCs), and fibro-adipogenic progenitors (FAPs) | Moiseeva et al., Nature, 2023 |
| Reyes_XDPI_UP_Lung_Injury | Mouse | No | 3 | in vivo | Fibroblasts | Reyes et al., Science, 2022 |
| EC.SENESCENCE.SIG | Human | No | 1 | Analysis of cancer scRNA-seq datasets | ECs in multiple cancer types | Wu et al., J Biomed Sci, 2023 |
| Aggregated SnG module | Human | Yes | 1 | Analysis of bulk RNA-seq of 50 tissues from GTEx | Multiple tissues | Xu et al., Molec Neurodegener, 2022 |
| CellAge | Human | No | 1 | Curated from literature | Several cell types | Avelar et al., Genome Biol, 2020 |
| SnC vs Quiescent Fibroblasts | Mouse | Yes | 2 | in vitro | Fibroblasts | Cherry et al., GeroScience, 2023 |
| SenSkin | Human | No | 1 | Curated from literature | Skin | Wyles et al., GeroScience, 2025 |
| CoreScence | Human | No | 1 | Curated from literature | N/A | Qu et al., Cell Genomics, 2025 |
| SenCat | Human | Yes | 1 | in vitro | N/A | Anerillas et al., Molecular Cell, 2026 |
| cogaps_k(18, 12)_senescence_fibrosis | Mouse | Yes | 12 and 9 | in vivo | Fibroblasts | Rindone et al., bioRxiv, 2026 |
| YE_BELLE_SENCAF | Mouse | Yes | 1 | in vivo | Fibroblasts | Ye et al., Cancer Discov, 2024<br>Belle et al., Cancer Discov, 2024 |
| SenSig | Mouse | Yes | 1 | in vivo | Fibroblasts | Cherry et al., Geroscience, 2023<br>Rindone et al., bioRxiv, 2026 |
| SenePy (Human) | Mouse | Yes and No | 75 and 1 | Analysis of *Tabula Muris Senis* scRNA-seq atlas | Multiple cell and tissue types | Sanborn et al., Nature Comm, 2025 |
| SenePy (Mouse) | Human | Yes and No | 65 and 1 | Analysis of scRNA-seq datasets from various tissue and cell types across human lifespan | Multiple cell and tissue types | Sanborn et al., Nature Comm, 2025 |

**Supplementary Table 2.** List of senescence signatures in SENgEAR.

| Treatment/Disease | Species | Tissue | Datatype | Reference |
|---|---|---|---|---|
| Keloid | Human | Skin | scRNA-seq | Deng CC et al., Nat Comm, 2021 |
| BCC | Human | Skin | scRNA-seq | Ganier C et al., Proc Natl Acad Sci USA, 2024 |
| IPF | Human | Lung | scRNA-seq | Habermann AC et al., Science Advances, 2020 |
| Bleomycin-induced injury | Mouse | Lung | scRNA-seq | Tsukui T, Wolters PJ, Sheppard D, Nature, 2024 |
| IPF and Bleomycin-induced fibrosis | Human & Mouse | Lung | Visium | Franzen L et al., Nature Genetics, 2024 |
| Young, aged mice w/wo genetic mutation that causes fibrosis | Mouse | Lung | scRNA-seq | Banaschewski BJH et al., Comm Biol, 2025 |
| PDAC | Human | Pancreas | scRNA-seq | Loveless IM et al., Clin Cancer Res, 2025 |
| Normal, NASH diet (liver only) | Mouse | Liver & Kidney | scRNA-seq | Omori S et al., Cell Metabolism, 2020 |
| Muscle injury treated with saline or biomaterial implants | Mouse | Skeletal muscle | scRNA-seq | Cherry C et al., GeroScience, 2023 |
| Foreign body response to synthetic implant in muscle injury | Mouse | Skeletal muscle | scRNA-seq | Ruta A et al.,Nature Communications, 2026 |
| Young, aged mice with myotoxin injury | Mouse | Skeletal muscle | scRNA-seq | Walter L et al., Nature Aging, 2024 |
| Sorted GFP+/- ( p21+/-) from aged mice | Mouse | Adipose | scRNA-seq | Weng B et al., Cell Metabolism, 2024 |
| Breast cancer | Human | Breast | scRNA-seq | Wu SZ et al., Nature Genetics, 2021 |

**Supplementary Table 3.** Analysis settings used for data visualizations. Links to each analysis are provided in Supplementary File 1.

| Dataset | Signature | Projection method | z-score normalization |
|---|---|---|---|
| Weng B et al., Cell Metabolism, 2024 | SenSig | ProjectR | Yes |
|  | SenMayo | PCA | No |
|  | SenCat | ProjectR | Yes |
| Habermann AC et al., Science Advances, 2020;Franzen L et al., Nature Genetics, 2024 | SenSig | ProjectR | Yes |
|  | SenMayo | PCA | No |
|  | SenCat | ProjectR | Yes |
|  | cogaps_k12_senescence_fibrosis | ProjectR | No |
| Ruta A et al.,Nature Communications, 2026 | SenSig | ProjectR | Yes |
|  | SenMayo | PCA | No |
|  | SenCat | ProjectR | Yes |
| Walter L et al., Nature Aging, 2024 | SenSig | ProjectR | Yes |
| Figure 1 and Cdkn2a expression | N/A | N/A | N/A |

